# Evo-Scope: Fully automated assessment of correlated evolution on phylogenetic trees

**DOI:** 10.1101/2022.12.08.519595

**Authors:** Maxime Godfroid, Charles Coluzzi, Amaury Lambert, Philippe Glaser, Eduardo P. C. Rocha, Guillaume Achaz

**Author notes:** Corresponding author: Maxime Godfroid.

## Abstract

1. Correlated evolution describes how multiple biological traits evolve together. Recently developed methods provide increasingly detailed results of correlated evolution, sometimes at elevated computational costs.
2. Here, we present ***evo-scope***, a fast and fully-automated pipeline with minimal input requirements to compute correlation between discrete traits evolving on a phylogenetic tree. Notably, we improve two of our previously developed tools that efficiently compute statistics of correlated evolution to characterize the nature, such as synergy or antagonism, and the strength of the interdependence between the traits.
3. Furthermore, we improved the running time and implemented several additional features, such as genetic mapping, Bayesian Markov Chain Monte Carlo estimation, consideration of missing data and phylogenetic uncertainty.
4. As an application, we scan a published penicillin resistance data set of *Streptococcus pneumoniae* and characterize genetic mutations that correlate with antibiotic resistance. The pipeline is accessible both as a self-contained github repository (https://github.com/Maxime5G/EvoScope) and through a graphical galaxy interface (https://galaxy.pasteur.fr/u/maximeg/w/evoscope).

## Introduction

The study of correlated evolution between biological traits illustrates the tight interdependence between processes in evolutionary biology. The constant development of new tools (listed in Table S1) to understand these dependencies now allow one to predict residues in contact in 3D structures of proteins (Morcos et al., 2011) or to identify protein-protein interactions (Barker & Pagel, 2005). Nevertheless, the measuring of these correlations is complicated by the phylogenetic relations (i.e., phylogenetic inertia) between living entities, typically represented by a phylogenetic tree. As such, species belonging to the same taxon may share several residues or biological traits by vertical descent, a process which one usually wants to disentangle from converge due to other processes, such as natural selection (Achaz & Dutheil, 2021). Therefore, the measurement of correlation between traits must either correct, or explicitly account for phylogenetic structure.

A typical application of the study of correlated evolution is the measure of the correlation between genetic factors and a given phenotype, i.e., genetic mapping, generally performed through genome-wide association studies (GWAS). Multiple tools have been devoted to measure such correlation, and these use different methods to account for phylogenetic relationships (see Table S1). While most methods add a correction term for this, few rely explicitly on the topology of the phylogenetic tree (e.g., treeWAS (Collins & Didelot, 2018)). Particularly in bacteria, tree-aware GWAS has been critical to detect genetic variants involved in the evolution of antibiotic resistance (Farhat et al., 2019), virulence (Galardini et al., 2020) or niche adaptation (Gori et al., 2020).

We have previously developed two methods to measure correlated evolution on phylogenetic trees. First, *epics* calculates all possible orderings of mutational events affecting two traits occurring on a phylogenetic tree, and computes efficiently the exact probability to observe a number of co-counts equal or larger than the observed repartition (Behdenna et al., 2016). The second tool, *epocs*, computes the influence of the mutation of a first trait on the mutation rate of a second one by maximum likelihood (Behdenna et al., 2021). Notably, *epocs* assesses different possible interactions between the traits, such as induction or inhibition.

Here we describe a novel software, *evo-scope* (EVOlutionary Study of Correlations of Occurrences on a Phylogeny), combining *epics* and *epocs*, to ease the detection of correlated evolution on phylogenetic trees for discrete traits. We further detail new features that we have added in the *evo-scope* implementation. Notable improvements include accelerated run-times (e.g., using sparse matrices in *epics*), consideration of missing data in species traits, analysis of a single trait (i.e., GWAS), the analysis of forests of trees and a Bayesian Markov Chain Monte Carlo (MCMC) estimation.

### Presentation of *evo-scope*

The details of the *epics* and *epocs* models are described in Behdenna (2016, 2021). *Evo-scope* provides new versions of these tools with additional features.

### Description of the pipeline

The *evo-scope* pipeline consists of five steps: ancestral character reconstruction, parsing and formatting, running *epics*, running *epocs* and summarizing the results (Fig 1).

**Figure 1.**
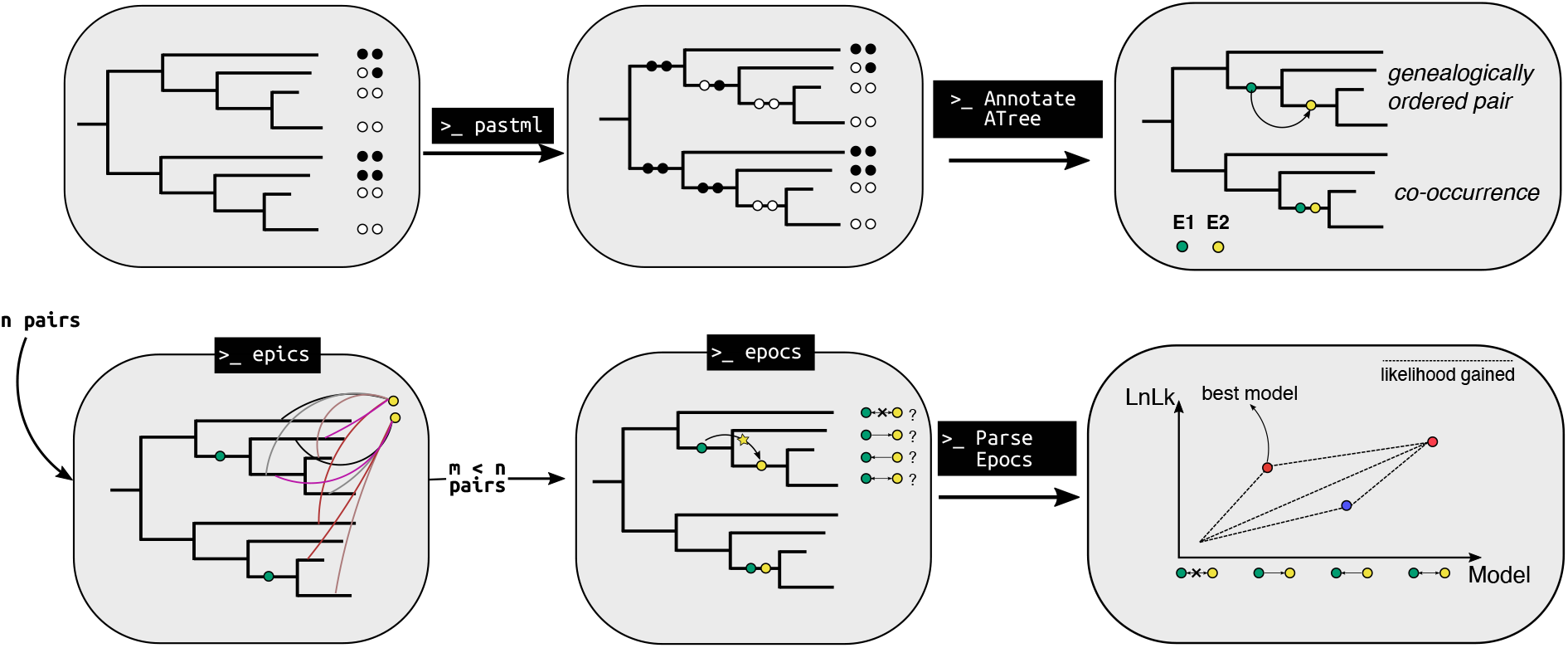
Schematic representation of the *evo-scope* pipeline (see Methods). The user supplies a rooted phylogenetic tree with the values of *n* discrete traits at the tips. Pastml reconstructs the evolution of each trait along the tree and an R script formats the reconstruction to display the mutations on the branches. *Epics* is run as a pre-filter to identify significantly associated pairs of evolutionary processes (*m*) among the initial *n*. Next, *epocs* maximizes the likelihoods of four evolutionary models, one of independence and three of dependence, on the selected pairs. Finally, an R script summarizes the results and outputs the best model explaining the repartition of events on the tree. Additionally, a figure can be generated for any pair considered to describe the gains of likelihood between nested models.

#### 1. Ancestral Character Reconstruction (ACR)

As initial input, a user provides a rooted phylogenetic tree and a file with the values of discrete traits at the tips. Pastml reconstructs the trait states on the internal branches of the tree using the JOINT model (Ishikawa et al., 2019; Pupko et al., 2000). Any other ACR tool or algorithm (e.g., Maximum A Posteriori in *pastml*) can be used as long as one single state (defined or undefined) is provided on each branch.

#### 2. ACR parsing and tree reformatting

*Evo-scope* reconstructs all mutational events on the branches based on the ancestral reconstruction. A mutational event is defined as any change of a trait value between two consecutive nodes of the tree. At most one mutation per branch per trait is allowed: either a change occurred between the parent and daughter node or none.

#### 3. *Epics* as a pre-filter

*Evo-scope* then runs *epics* to discover by default pairs of traits that co-occur frequently on the branches of the trees (Behdenna et al., 2016). Other scenarios for the pairs of traits placement are available, such as genealogically ordered pairs. At this step, *epics* is used as a pre-filter as it runs very fast and it only detects statistical association between the mutational events on the tree. Importantly, *epics* does not allow to predict an evolutionary scenario. As such, additional in-depth analysis is required to better describe the interactions between the mutational events. However, for very large data sets, *epics* might be the only option available as running time increase substantially for the maximum likelihood-based tool *epocs*. The output of this step is the list of all pair of events that are significantly associated on the tree for a given p-value.

#### 4. *Epocs* for the model discovery

For all significantly associated traits, *epocs* maximizes the likelihood of multiple scenarios of induction to determine which of the trait influenced the occurrence of the other one. The multiple scenarios can contain from two to eight parameters and describe the interrelationship between the occurrence of two events on the tree. The parameters are divided into natural occurrence rates (*μ*_{*i,j*}_, υ_{*i,j*}_) and excited occurrence rates 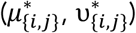 for each of the trait (Fig 3A). The ratio between the excited occurrence rates and the natural occurrence rates defines the induction.

In the current version, the model exploration is restricted to three models of correlation compared to a model of independence. The three models of correlation describe either an induction of the first trait on the second one, vice-versa or a co-induction of both events.

#### 5. Summarizing the results

Finally, *evo-scope* extracts the likelihoods and parameter values inferred for each of the models. To select the best model for each pair, the program performs likelihood ratio tests (LRT) between nested models (Behdenna et al., 2021). Next, it compares the calculated LRT value to a *χ*^2^ distribution whose degrees of freedom is the difference in the number of parameters between models. Finally, based on significant LRTs, *evo-scope* selects the “best” model by a trade-off measure which maximizes the LRT value and minimizes the number of model parameters. *Evo-scope* then allows the user to select a pair and plot the likelihood gains between nested models.

### Additional novel features to *epics* and *epocs*

#### 1. Analysis on multiple independent trees

The user can provide a list of independent trees (non-overlapping taxa), to either *epics* or *epocs*, with the same type of events. This feature is particularly useful whenever the same character can be analyzed across unrelated clades. In particular, *epics* and *epocs* will collect the information of all the trees and generate a global result by integrating the information across all trees. To do so, *epics* computes an aggregated p-value convoluted across all trees, while *epocs* transmits the likelihood calculation across all trees.

#### 2. Forest of trees

To account for phylogenetic uncertainty, the user can provide a forest of trees, such as generated by BEAST or MrBayes (Ronquist et al., 2012; Suchard et al., 2018), to either *epics* or *epocs*, and the results are ranked to extract the mode and 95% credibility intervals. In *epics*, the p-values are ranked by the tree likelihoods, while in *epocs* the results are ranked by the product of the *epocs* likelihood and the tree likelihood.

#### 3. MCMC exploration of likelihood surface

In the standard *epocs*, the likelihood is maximized considering all possible arrangements, with some restrictions on the number of co-occurrences due to computational complexity. In the MCMC case, the program is allowed to explore freely any possible arrangement of the co-occurrences. This feature is implemented in a third executable *epocs_mcmc*. In this module, the user supplies a tree with events to analyze and a standard Metropolis-Hastings algorithm will explore the parameter distribution across the number of replicates selected by the user (Hastings, 1970). Notable behaviors of the MCMC chain determine the relevancy of each parameter in the model. For example, a flat uniform distribution of the posterior is likely to reflect that this parameter is not pertinent in the model. In addition, the algorithm explores the order in the co-occurrences of the traits. *Epocs_mcmc* outputs a tab-delimited file compatible with Tracer (Rambaut et al., 2018).

#### 4. Missing data

The presence of uncharacterized traits at the tips can be taken into account in each of the tools above. For *epics*, the branches where traits are unknown are masked, whereas for *epocs* and *epocs_mcmc*, branches containing an unknown trait value and those below are excluded in the likelihood calculation.

#### 5. GWAS-like procedure

*Epics* and *epocs* have now the possibility of selecting either one character or two to test in the procedures. Whenever the user selects only one character, the analysis is similar to a GWAS, where any other character is tested against the one selected.

### Analysis of an antibiotic resistance dataset

As test data, we retrieved 603 genomes of *Streptococcus pneumoniae* (Chewapreecha et al., 2014; Croucher et al., 2013, 2015; Lees et al., 2018). In this data set, the resistance to penicillin is known and encoded as a binary variable (R for resistant strains, S for susceptible strains). Also, 198,248 single-nucleotide polymorphisms (SNPs) have been called in the genomes. For the post-hoc MCMC analysis, we retrieved a significant association from *evo-scope* and ran *epocs_mcmc* with a chain length of 10^7 steps and sampling every 50 steps on this pair.

### Availability

The source code of *evo-scope, epics* and *epocs* are available on github under the GNU General Public License v3: https://github.com/Maxime5G/EvoScope, and pastml is available either on bioconda (Grüning et al., 2018) or on github: https://github.com/evolbioinfo/pastml. We also provide a galaxy access to all tools and a “push-button” workflow at the Pasteur institute server: https://galaxy.pasteur.fr/u/maximeg/w/evoscope.

## Results

To illustrate the applicability of the present pipeline, we evaluated *evo-scope* by performing a GWAS-like analysis to a data set of 603 genomes of *S. pneumoniae* (Chewapreecha et al., 2014; Croucher et al., 2013, 2015; Lees et al., 2018). As a positive control, *evo-scope* detected genetic variants correlated to penicillin resistance (following (Chewapreecha et al., 2014)). In addition, *evo-scope* inferred the induction values for the significant associations to determine the strength and the direction of the correlation between the occurrence of antibiotic resistance and the SNPs. Of the 82,829 SNPs present in at least eight genomes, we report a total of 1,629 SNPs (1.97%) associated with the resistance by *epics* (Fig. 2A) – of which 543 (33.3% of the significant SNPs) were further inferred to be correlated with the resistance by *epocs*, with a P-value < 10^−4^ (Fig. 2B,C). In the end, *evo-scope* found that nearly half of the correlated SNPs (n=255, 47%) are located in only three genes, *penA, pbpX* and *pbp1A*, in accordance with previous results, as mutations in these three genes are the major drivers of penicillin resistance in *S. pneumoniae* (Chewapreecha et al., 2014) (Fig. 2B,C). In addition, we detected 45 significant SNPs in *mraY*, a gene involved in cell wall biogenesis (Chewapreecha et al., 2014), and 44 SNPs in *clpL*, reported to be related to penicillin susceptibility and compensation of the antibiotic resistance fitness cost (Hakenbeck et al., 2012; Tran et al., 2011). In our study, most of the inferred inductions revealed that the resistance phenotype occurred first in the tree, with the genetic mutations occurring very quickly afterwards (i.e., induction values > 100 – Fig 2C).

**Figure 2.**
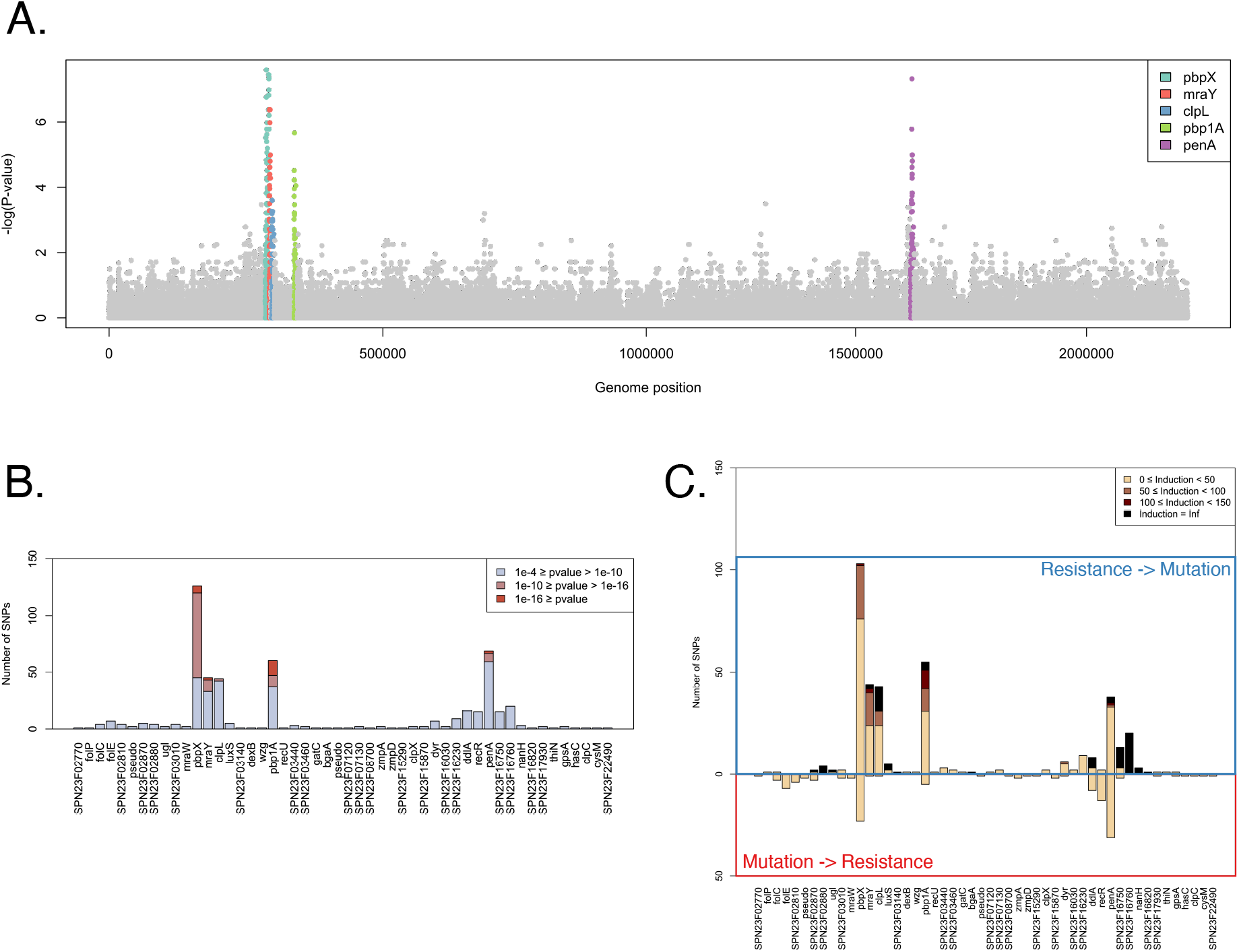
Application of *evo-scope* to a dataset of *S. pneumoniae*. (A) Manhattan plot of the P-values inferred by *epics*. In color are shown genes of interest with multiple SNPs of low p-values. Notably, the three highest peaks correspond to *pbpX, pbp1A* and *penA*. (B) Barplot showing the p-value inferred by *epocs*, grouped by gene IDs. (C) Barplots showing the inductions inferred by *epocs* (*i*.*e*., the lambda in (Behdenna et al., 2021)). Top row represents the inductions Resistance -> Mutation, bottom row represents the other way round. For each statistical association between the resistance and one of the 543 significant SNP, we retrieved the best model using LRT. Most notably, the three genes of interest *pbpX, penA* and *pbp1A* display low P-values and strong inductions.

Additionally, we include a post-hoc MCMC analysis with *epocs_mcmc* to demonstrate the usefulness of this addition in the *evo-scope* toolbox. For this purpose, we tested it on a significant association from *evo-scope*, which indicated an induction resistance -> mutation. (Table 1 and Fig. 3B).

**Table 1.** Parameter estimates of the MCMC run for the selected pair.

| *Parameter* | *ML estimates* | *Mean of the bayesian estimates* | *Mode of the posterior distribution* | *95% Credible interval* |
| --- | --- | --- | --- | --- |
| $\mu 1$ | 46.263 | 41.03 | 38.81 | [25.39 – 57.51] |
| $\mu 1^*$ | 292.22 | 543.25 | 503.8 | [214.42 – 934.32] |
| $\mu 2$ | 6.8197 | 14.66 | 12.79 | [4.41 – 25.73] |
| $\mu 2^*$ | 377.08 | 338.24 | 317.8 | [148.78 – 543.70] |
| $\nu 1$ | 41.474 | 46.23 | 41.76 | [20.36 – 73.91] |
| $\nu 1^*$ | ? | 475.22 | 68.12 | [0.33 – 37.86] |
| $\nu 2$ | 0 | 4.67 | 0.5724 | [1.4E-5 – 13.94] |
| $\nu 2^*$ | 0 | 294.33 | 41.59 | [3.6E-5 – 41.23] |

**Figure 3.**
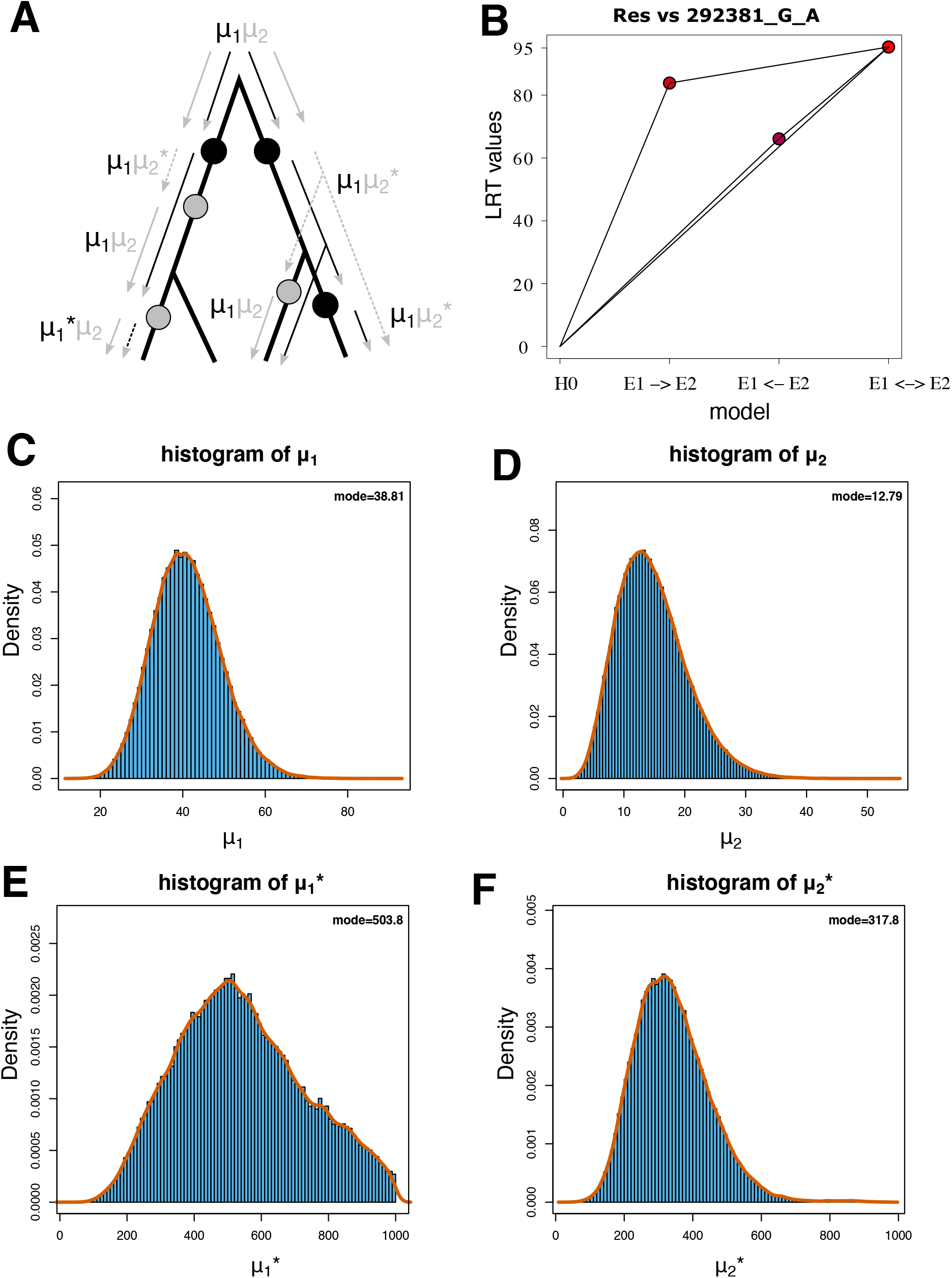
Likelihood ratios and output from an *epocs_mcmc* run on one significant association between the resistance phenotype and a SNP in *pbpX*. (A) Summary graph of parameter models and transitions upon evolutionary events on a mock tree, adapted from (Behdenna et al., 2021). Non-starred rates are natural rates of occurrence, starred rates are excited rates of occurrence. An occurrence of the first event in the pair activates the excited rates of the second event in the pair. Once the second event occurs, the excitation is consumed. (B) Likelihood ratios between the four models tested by *evo-scope*, following the convention from (Behdenna et al., 2021). Lines connect nested models. The comparison providing the best likelihood gain while minimizing the number of parameters is the comparison between the model of independence and the model of induction E1->E2 (i.e., Resistance -> Mutation). (C-F) Histograms of parameter values taken from the MCMC chain. Scales are parameter-based in order to best display the shape of the parameter distribution.

After running *epocs_mcmc*, we see that ***μ***1 is clearly defined, with a narrow distribution (Fig 3C). Interestingly, even though the preferred model by ML is the induction Resistance -> Mutation, ***μ***1* (demonstrating an induction Mutation -> Resistance) is still somewhat relevant to the model, but the confidence interval is quite wide (Fig 3E). Importantly, ***μ***2 is close to zero and ***μ***2* is clearly defined with a narrow distribution (Fig 3D,F). We conclude from these observations that the selected mutation in *pbpX* mostly occurs after the resistance, therefore indicating an induction Resistance -> Mutation. In addition, the order of the co-occurrences are inferred in the same direction as ML and further, the **υ** parameter posterior distributions are centered around the parameter values inferred by ML (Fig S1).

## Discussion

Inferring correlations between evolutionary events has multiple useful applications. However, fast and easily interpretable tools are still lacking. Here, we provide a self-contained and fully automated pipeline to detect correlated evolution, with minimal input required from the user. *Evo-scope* takes as input any number of discrete traits and a rooted phylogenetic tree and outputs a model of correlated evolution best explaining the repartition of the events on the tree. The flexibility of *evo-scope* allows to perform different types of analyses, such as GWAS as exemplified in this paper. Here, we applied *evo-scope* on a well-described dataset of antibiotic resistance and SNPs in *S. pneumoniae* to 1) test the accuracy of the pipeline and 2) to add on the literature possible inductions either from the mutations to the resistance or in the other way round. We showed that *evo-scope* is able to retrieve consistent results with the literature in a timely manner.

The Induction values inferred on the significantly associated pairs revealed that most of the mutations associated with the resistance occurred after the resistance phenotype. Two effects may contribute to explain this result. Methodologically, the ACR step might introduce biases regarding the real occurrence timing of both mutations and resistance as the algorithm tries to minimize the number of steps to reconstruct a trait, while antibiotic resistance and resistance-conferring mutations are known to be subject to homoplasies (see for example (Coolen et al., 2021)).

In bacteria, genomic changes that are necessary to evolve antibiotic resistance are often a burden for the bacteria, imposing a fitness cost (Andersson & Hughes, 2010). However, mechanisms exist that compensate the cost of such antibiotic resistance, such as the evolution of a secondary mutation in the genome. For *S. pneumoniae*, bacteria exhibiting multiple mutations in the penicillin binding proteins exhibit a phenotype of resistance and compensation (Orio et al., 2011). Using *evo-scope*, we showed that many significantly associated pairs of mutations concur with these observations. Furthermore, the patterns of mutation acquisitions and inductions could shed light on evolutionary trajectories of the acquisition of antibiotic resistance in *S. pneumoniae*.

In conclusion, we developed a fully-automated pipeline, with minimal input required from the user. Our approach, by considering explicitly the phylogenetic component, allows to detect correlated evolution between discrete traits of any species by explicitly taking into account the underlying structure. We show that our pipeline can also be applied to tree-aware GWAS analyses in bacteria, expanding previous results in the field linking genetic variants to penicillin resistance in *S. pneumoniae*.

## Acknowledgements

This work used the computation and storage services (TARS cluster) provided by the IT department at Institut Pasteur, Paris.

## Author contribution

GA, MG, AL, ER, PG, CC conceived the ideas and designed methodology; MG collected the data; GA, MG analyzed the data; MG led the writing of the manuscript. GA, AL, PG, ER obtained the funding for this project. All authors contributed critically to the drafts and gave final approval for publication.

