## Supplementary material for "Evo-Scope: Fully automated assessment of correlated evolution on phylogenetic trees": Fig S1

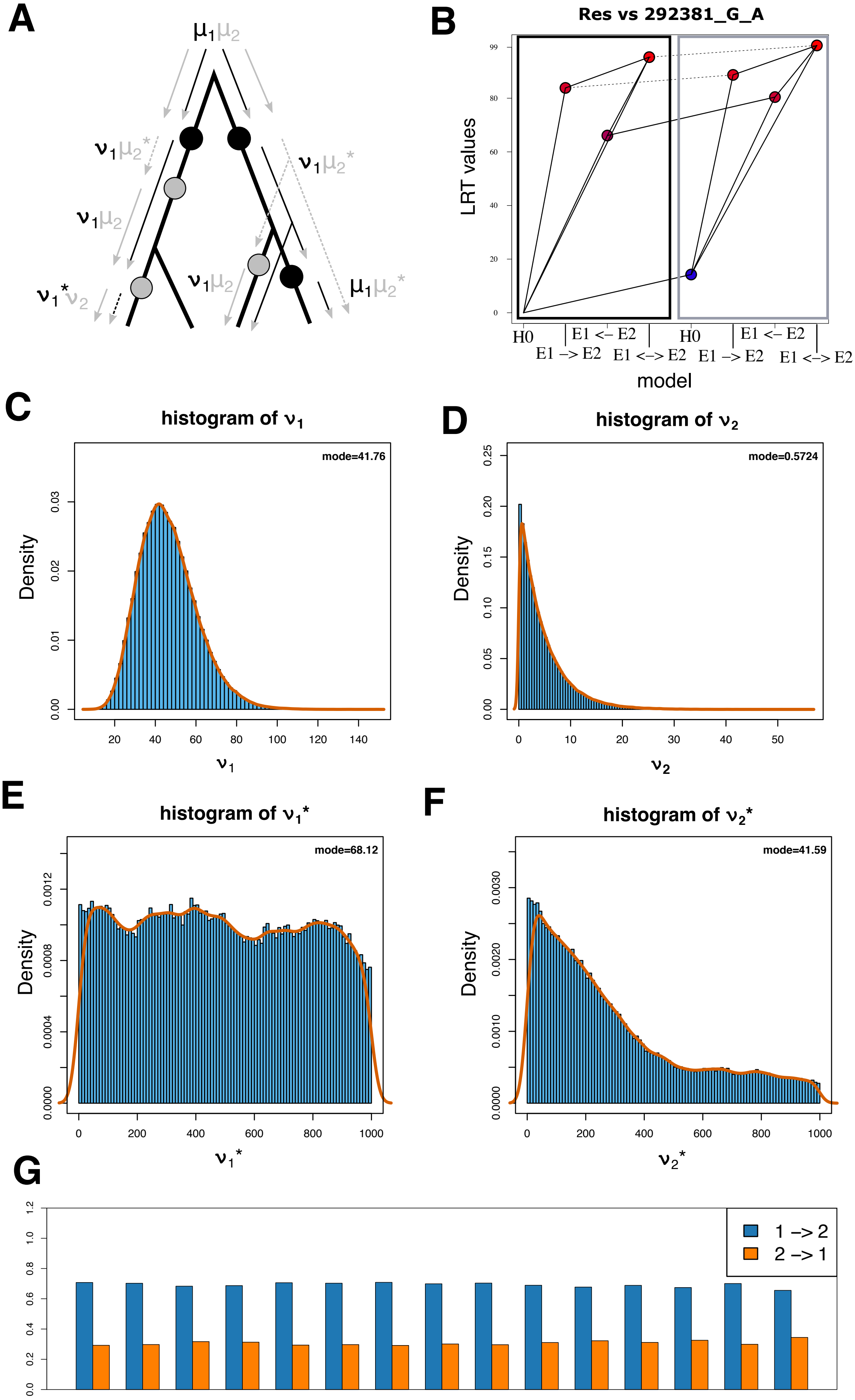

A
B
Res vs 292381_G_A
LRT values
H0
E1 -> E2
E1 <- E2
E1 <-> E2
model
C
histogram of ν1
mode=41.76
Density
ν1
D
histogram of ν2
mode=0.5724
Density
ν2
E
histogram of ν1*
mode=68.12
Density
ν1*
F
histogram of ν2*
mode=41.59
Density
ν2*
G
1 -> 2
2 -> 1

**Figure S1. Likelihood ratios and output from epocs_mcmc, with the state-dependent parameters, on one significant association between the resistance phenotype and a SNP in *pbpX*.** (A) Summary graph of parameter models and transitions upon evolutionary events on a mock tree, adapted from (Behdenna et al., 2021). Non-starred rates are natural rates of occurrence, starred rates are excited rates of occurrence. An occurrence of the first event in the pair activates the excited rates of the second event in the pair. Once the second event occurs, the excitation is consumed. (B) Likelihood ratios between the eight models tested by scoop. Lines connect nested models, where plain lines show significant LRT and dotted lines show non-significant LRT. The black square shows the LRTs for models without state-dependence and the light grey square shows the LRTs for the models with state-dependence. The comparison providing the best likelihood gain while minimizing the number of parameters is the comparison between the model of independence and the model of induction E1->E2 (i.e., Resistance -> Mutation) with state-dependence, (C)-(F) histogram of parameter values taken from the MCMC chain for the state-dependence parameters, (G) order of mutations in the co-occurrences in the branches of the tree. We observed that the MCMC algorithm selected preferentially the direction E1 -> E2 (i.e., Resistance -> Mutation) between 65.5% (co-occurrence 15) and 70.8% (co-occurrence 7) of the time.
